# The role of metals in hypothiocyanite resistance in *Escherichia coli*

**DOI:** 10.1101/2024.03.07.583962

**Authors:** Michael J. Gray

**Author notes:** Address correspondence to Michael J. Gray.

## Abstract

The innate immune system employs a variety of antimicrobial oxidants to control and kill host-associated bacteria. Hypothiocyanite/hypothiocyanous acid (^-^OSCN/HOSCN) is one such antimicrobial oxidant that is synthesized by lactoperoxidase, myeloperoxidase, and eosinophil peroxidase at sites throughout the human body. HOSCN has potent antibacterial activity while being largely non-toxic towards human cells. The molecular mechanisms by which bacteria sense and defend themselves against HOSCN have only recently begun to be elaborated, notably by the discovery of bacterial HOSCN reductase (RclA), an HOSCN-degrading enzyme widely conserved among bacteria that live on epithelial surfaces. In this paper, I show that Ni^2+^ sensitizes *Escherichia coli* to HOSCN by inhibiting glutathione reductase, and that inorganic polyphosphate protects *E. coli* against this effect, probably by chelating Ni^2+^ ions. I also found that RclA is very sensitive to inhibition by Cu^2+^ and Zn^2+^, metals that are accumulated to high levels by innate immune cells, and that, surprisingly, thioredoxin and thioredoxin reductase are not involved in HOSCN stress resistance in *E. coli*. These results advance our understanding of the contribution of different oxidative stress response and redox buffering pathways to HOSCN resistance in *E. coli* and illustrate important interactions between metal ions and the enzymes bacteria use to defend themselves against oxidative stress.

**IMPORTANCE:** Hypothiocyanite (HOSCN) is an antimicrobial oxidant produced by the innate immune system. The molecular mechanisms by which host-associated bacteria defend themselves against HOSCN have only recently begun to be understood. The results in this paper are significant because they show that the redox buffer glutathione and enzyme glutathione reductase are critical components of the *Escherichia coli* HOSCN response, working by a mechanism distinct from that of the HOSCN-specific defenses provided by the RclA, RclB, and RclC proteins, and that metal ions (including nickel, copper, and zinc) may impact the ability of bacteria to resist HOSCN by inhibiting specific defensive enzymes (*e.g.* glutathione reductase or RclA).

## INTRODUCTION

Hypothiocyanite/hypothiocyanous acid (^-^OSCN/HOSCN) is an antimicrobial pseudohypohalous acid synthesized from hydrogen peroxide (H_2_O_2_) and thiocyanate (SCN^-^) by mammalian heme peroxidases, including lactoperoxidase (LPO), myeloperoxidase (MPO), and eosinophil peroxidase (EPO)(1, 2). HOSCN is less reactive than the other major antimicrobial products of heme peroxidases, particularly when compared to the extremely reactive hypochlorous acid (HOCl) produced by MPO (1, 3). HOSCN is a specific thiol-oxidizing agent (2, 3) and is essentially non-toxic to human cells, a fact which has been attributed to the ability of mammalian selenocysteine-containing thioredoxin reductase to reduce HOSCN to nontoxic SCN^-^ and H_2_O (4). In contrast, bacterial thioredoxin reductase is inhibited by HOSCN (4), as are a number of central bacterial metabolic enzymes (5–7). HOSCN has therefore long been considered a specifically antimicrobial product which the innate immune system uses to control bacteria without causing damage to host tissues, a model which has generally assumed that bacteria lack effective defenses against HOSCN (8–11).

That assumption has recently been shown to be an oversimplification, as recent studies from a number of laboratories have identified HOSCN-specific stress responses in a variety of host-associated bacteria (11). We identified RclA as an HOSCN-induced, highly-active bacterial HOSCN reductase that protects *Escherichia coli* and some other intestinal bacteria from HOSCN stress (12), followed rapidly by other research groups showing that the RclA homologs from *Streptococcus pneumoniae* (Har) and *Staphylococcus aureus* (MerA) played similar roles in those species (13, 14). *Pseudomonas aeruginosa* lacks an RclA homolog, but mounts an HOSCN stress response that depends on inorganic polyphosphate and a peroxiredoxin-like protein of unknown function called RclX (15, 16). The HOSCN stress response of *S. pneumoniae* is perhaps the best characterized among these bacteria, and studies in that species have identified both HOSCN-specific defense mechanisms (*i.e.* HOSCN reductase) and overlaps with more general oxidative stress and redox homeostasis mechanisms (*e.g.* the glutathione and thioredoxin systems)(14, 17–21).

Older studies of the antimicrobial activity of LPO generally did not use purified HOSCN, as we and the other labs cited in the previous paragraph have done more recently. Instead, they typically added a combination of LPO, SCN^-^, and one of a variety of H_2_O_2_-generating enzymatic systems to synthesize HOSCN *in situ* (22–25). This is in some ways more representative of the *in vivo* conditions in which bacteria are exposed to HOSCN, but makes it more difficult to control HOSCN concentrations or to separate bacterial responses to HOSCN from those to H_2_O_2_ or to other products of LPO (1, 26–29). Roughly 20 years ago, the laboratory of C.W. Michiels carried out a series of experiments to characterize the response of *E. coli* to the LPO / SCN^-^ / H_2_O_2_ system, in which they found that high pressure sensitizes *E. coli* to killing by this system (30, 31), that outer membrane lipids and porins play an important role in sensitivity (32), and that LPO / SCN^-^ / H_2_O_2_ induces a different response than other oxidative stressors (33, 34). Curiously, they also observed that exogenous nickel (Ni^2+^) sensitized *E. coli* to LPO / SCN^-^ / H_2_O_2_ (35), but provided no mechanistic explanation for this phenomenon.

In this paper, I confirm that Ni^2+^ does indeed sensitize *E. coli* to purified HOSCN and show that it does so by inhibiting glutathione reductase (Gor) activity. Since HOSCN efficiently inhibits *E. coli* thioredoxin reductase (TrxB)(4), this means that the combination of HOSCN and Ni^2+^ simultaneously disrupts both of *E. coli*’s cytoplasmic redox buffering systems, rendering it very sensitive to oxidative damage (36). I also show that RclA is subject to inhibition by metals, including Ni^2+^ but much more sensitively by Cu^2+^ and Zn^2+^. However, Ni^2+^-sensitization of *E. coli* to HOSCN was not RclA-dependent. These results confirm that low molecular weight thiols like glutathione are important conserved HOSCN resistance elements in bacteria (19, 37), help connect the older literature on bacterial responses to LPO to current work on the molecular mechanisms of HOSCN responses, and identify new connections between metal ions and the enzymes bacteria use to defend themselves against oxidative stress.

## RESULTS

### Nickel enhances HOSCN toxicity in *E. coli*

Sermon *et al*. (35) reported in 2005 that *E. coli* is sensitized to killing by the LPO / SCN^-^ / H_2_O_2_ system in the presence of 300 µM Ni^2+^, and that mutants lacking the Ni^2+^ / Co^2+^ / Mg^2+^ transport protein CorA (38, 39) are protected from this effect. They did not propose a mechanism to explain this unusual phenotype. Recent work from my lab and others (12–19) has now begun to identify the molecular mechanisms underlying HOSCN resistance in bacteria, and I therefore revisited this observation to see if I could gain new insights into how Ni^2+^ or other metal ions might impact that process. First, I tested whether Ni^2+^ could sensitize *E. coli* to growth inhibition by purified HOSCN in minimal medium, the method which has been used in most of the recent papers examining bacterial HOSCN response (12–14, 17–19), rather than by the more complex enzymatic LPO / SCN^-^ / H_2_O_2_ system (1, 11, 40). As shown in **Fig 1**, in this assay addition of 300 µM NiSO_4_ greatly enhanced the toxicity of 400 µM HOSCN for the wild-type, but not for the Δ*corA* mutant, consistent with the results of Sermon *et al.* (35). 300 µM NiSO_4_ did not inhibit growth of *E. coli* under these conditions (**Supplemental Fig S1A**) and neither did SCN^-^ at concentrations up to 40 mM (**Supplemental Fig S1B**). Addition of SCN^-^ did not change the minimal inhibitory concentration of NiSO_4_ (1.25 – 2.5 mM)(**Supplemental Fig S1C**) and Ni^2+^ did not react directly with HOSCN (**Supplemental Fig S2**), indicating that the phenotype seen in **Fig 1** was due to a physiological impact of Ni^2+^ on *E. coli* that sensitizes the cells to HOSCN.

**FIG 1.**
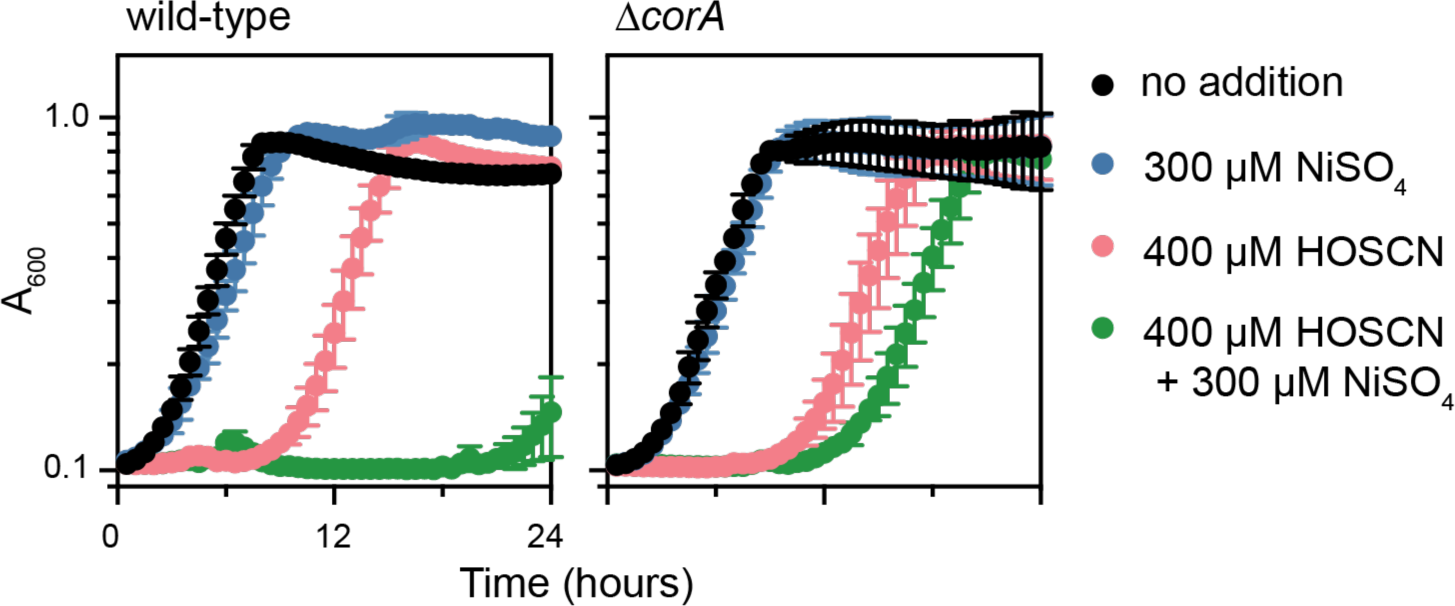
Nickel enhances HOSCN toxicity. *E. coli* strains MG1655 (wild-type) and MJG2314 (MG1655 &*corA761*::*kan*^+^) were grown in MOPS minimal medium without micronutrients containing 0.2% glucose, supplemented with 300 µM NiSO_4_ and/or 400 µM HOSCN, as indicated, incubating at 37°C with shaking and measuring A_600_ at 30-minute intervals for 24 hours (n=3 experimental replicates, with 3 technical replicates per experiment; error bars of 1 standard deviation).

### Inorganic polyphosphate protects *E. coli* against nickel-dependent HOSCN sensitization

Inorganic polyphosphate (polyP) is a bacterial stress response effector that protects against oxidative stress by chelating metal ions and stabilizing damaged proteins (41–43). Mutants of *Pseudomonas aeruginosa* lacking the *ppk1* gene encoding polyP kinase, which are therefore defective in polyP synthesis (44), are sensitive to inhibition by HOSCN (16). In contrast, *E. coli* Δ*ppk* mutants, which completely lack polyP (41, 45), were not more sensitive to HOSCN except in the presence of NiSO_4_ (**Fig 2**), indicating that the primary role of polyP in the *E. coli* HOSCN response, at least under these growth conditions, is to shield against the HOSCN toxicity-enhancing effect of metals. A mutant lacking *ppx*, encoding the polyP-degrading exopolyphosphatase PPX (46), was slightly more sensitive to HOSCN than the wild-type, but, like the wild-type, did not become more sensitive when NiSO_4_ was added under these conditions (**Fig 2**). It is worth noting that I performed the experiments in **Fig 2** in minimal medium supplemented with 0.1% casamino acids, since *ppk* mutants grow very poorly without amino acids (47, 48), and that the sensitivity of *E. coli* to both HOSCN and NiSO_4_ was somewhat different under these conditions from that seen in unsupplemented minimal medium (*e.g.* **Fig 1**).

**FIG 2.**
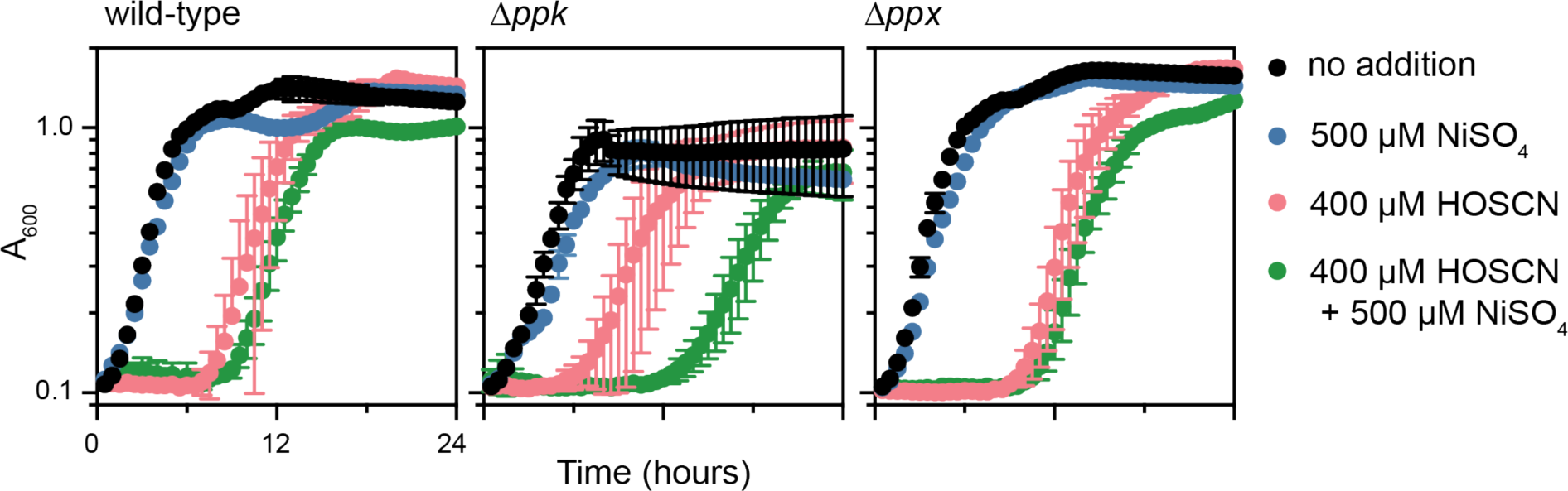
Polyphosphate protects *E. coli* against nickel-dependent HOSCN sensitization. *E. coli* strains MG1655 (wild-type), MJG0224 (MG1655 &*ppk-749*), and MJG0315 (MG1655 &*ppx-750*) were grown in MOPS minimal medium without micronutrients containing 0.2% glucose and 0.1% casamino acids, supplemented with 500 µM NiSO_4_ and/or 400 µM HOSCN, as indicated, incubating at 37°C with shaking and measuring A_600_ at 30-minute intervals for 24 hours (n=3 experimental replicates, with 4 technical replicates per experiment; error bars of 1 standard deviation).

### Copper, zinc, and nickel inhibit RclA activity *in vitro*

The *E.coli* HOSCN reductase RclA (12) (homologs of which are known as Har or MerA in *Streptococcus pneumoniae* and *Staphylococcus aureus*, respectively)(13, 14), is a flavin-dependent oxidoreductase that efficiently degrades HOSCN and protects bacterial cells against HOSCN toxicity, but is also known to interact with a variety of metal cations, notably Cu^2+^ and Hg^2+^ (49–51). One possible explanation for sensitization of *E. coli* to HOSCN by Ni^2+^, therefore, is that the metal ions might inhibit RclA’s HOSCN reductase activity, presumably by binding to the active site cysteine residues of RclA (50). *In vitro* assays with purified RclA (12, 49) and a variety of biologically relevant divalent metals revealed that Cu^2+^ and Zn^2+^ were extremely potent RclA inhibitors, capable of significantly inhibiting HOSCN reductase activity at concentrations as low as 1 µM (**Fig 3A**). Neither Cu^2+^ nor Zn^2+^ reacted with HOSCN on their own (**Supplemental Fig S2**). Ni^2+^ also significantly inhibited RclA activity, but only at much higher concentrations (>100 µM)(**Figs 3A,B**). Ca^2+^, Co^2+^, Mg^2+^, and Mn^2+^ had no impact on RclA activity (**Fig 3A**).

**FIG 3.**
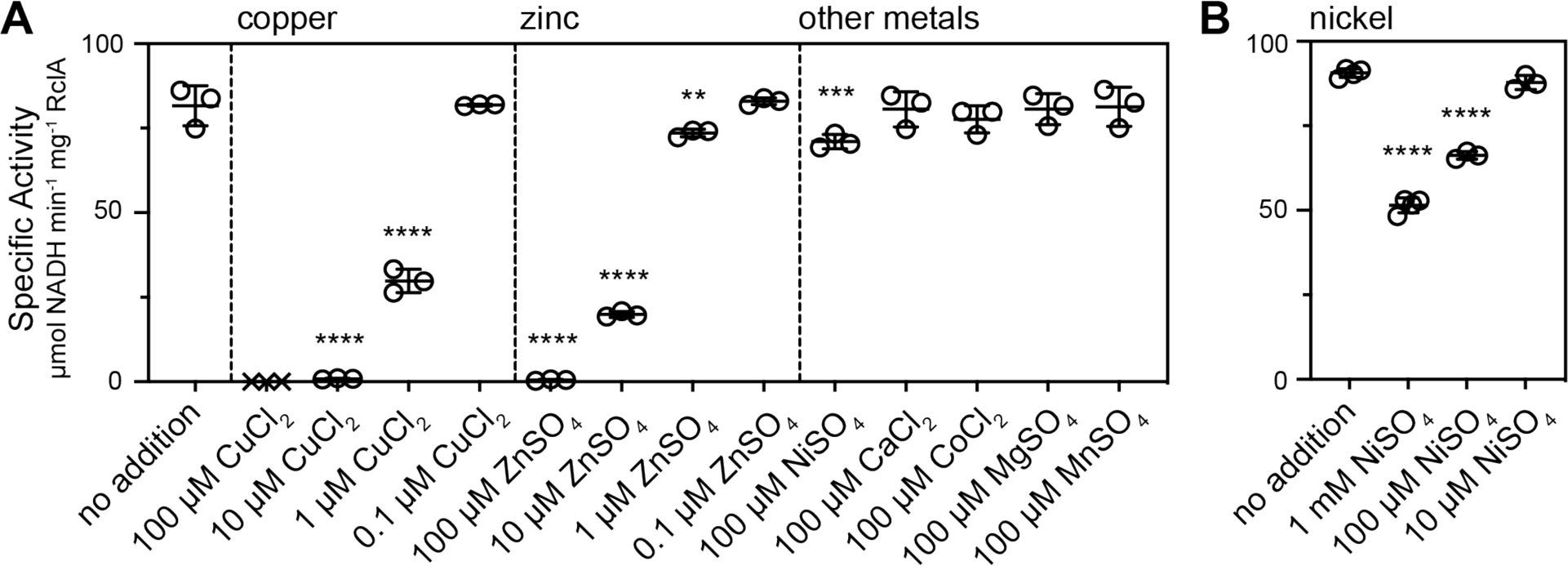
Copper, zinc, and nickel inhibit RclA activity *in vitro*. Reactions (500 µl) contained 10 mM HEPES-KOH buffer (pH 7.4), 160 µM NADH, 100 µM HOSCN, and the indicated concentrations of metal salts, and were started by addition of 10 nM RclA. I quantified NADH consumption at A_340_ for 1 min at 20°C and calculated specific activities as µmol NADH consumed min^-1^ mg^-1^ RclA (n=3-4 experimental replicates; error bars of 1 standard deviation). Asterisks indicate activities significantly different from that of RclA with no metals added in a given set of experiments (one-way ANOVA with Dunnett9s multiple comparisons test; ** = P<0.01. *** = P<0.001, **** = P<0.0001). X symbols indicate that RclA precipitated at 100 µM CuCl_2_.

### Sensitization of *E. coli* to HOSCN by nickel is independent of RclA, RclB, and RclC

Testing whether metals could potentiate HOSCN toxicity by inhibiting RclA *in vivo* presented some technical challenges. While NiSO_4_ was soluble in MOPS minimal medium up to a concentration of at least 40 mM (**Supplemental Fig S1A**), both CuCl_2_ and ZnSO_4_ formed visible precipitates (presumably insoluble phosphate salt crystals) when added to MOPS medium at concentrations higher than 1 µM (data not shown). At this concentration, CuCl_2_, the most potent inhibitor of RclA *in vitro* (**Fig 3A**), had no effect on the HOSCN sensitivity of either wild-type or Δ*rclA E. coli* (**Fig 4**). To try to address this, I deleted the *copA* gene, encoding the Cu export proteins CopA(Z) (38, 52, 53), which would be expected to increase the accumulation of Cu in the cytoplasm in *E. coli*. While this mutation did indeed confer a detectable Cu sensitivity phenotype at 1 µM CuCl_2_, it did not have a large impact on HOSCN sensitivity, and notably, any such impact was also apparent in the Δ*rclA* Δ*copA* double mutant (**Fig 4**), indicating that Cu was probably not affecting HOSCN sensitivity in this case by inhibiting RclA. I used a similar approach with the Zn and Ni exporters ZntA and RcnA (**Supplemental Fig S3**)(38, 54, 55), and found that deletion of *rcnA*, which did substantially sensitize *E. coli* to NiSO_4_ (**Supplemental Fig S4**), led to a small increase in HOSCN toxicity in the presence of 100 µM NiSO_4_, but not 1 µM CuCl_2_ or ZnSO_4_. None of the export mutants led to detectable increases in HOSCN sensitivity in the presence of 1 µM ZnSO_4_ (**Supplemental Fig S3**).

**FIG 4.**
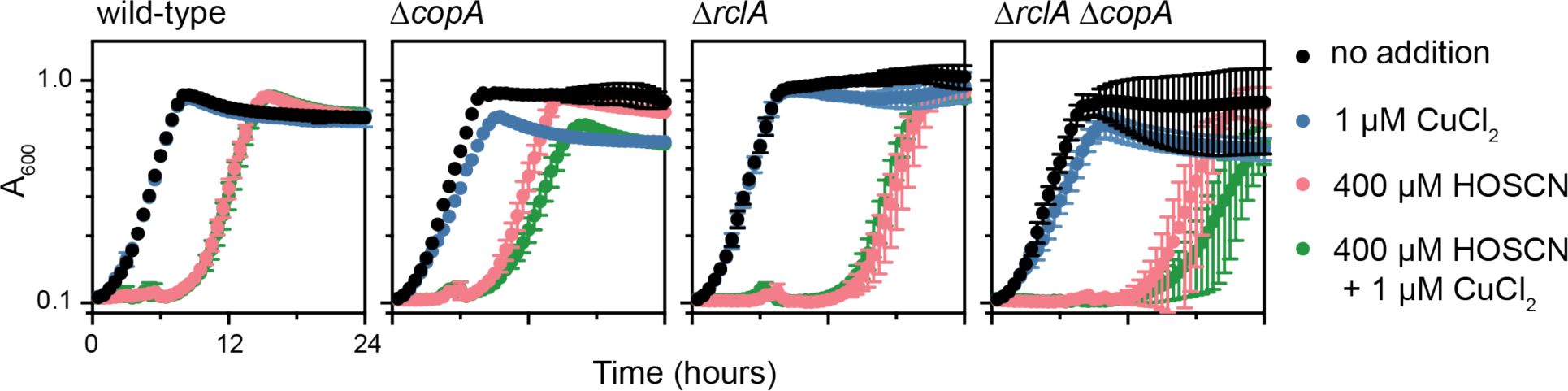
Sensitization of *E. coli* to HOSCN *in vivo* by copper is slight, and not RclA-dependent. *E. coli* strains MG1655 (wild-type), MJG1759 (MG1655 &*copA767*::*kan*^+^), MJG1958 (MG1655 &*rclA747*), and MJG2358 (MG1655 &*rclA747* &*copA767*::*kan*^+^) were grown in MOPS minimal medium without micronutrients containing 0.2% glucose, supplemented with 1 µM CuCl_2_ and/or 400 µM HOSCN, as indicated, incubating at 37°C with shaking and measuring A_600_ at 30-minute intervals for 24 hours (n=4 experimental replicates, with 3 technical replicates per experiment; error bars of 1 standard deviation).

*E. coli* Δ*rclA* mutants are more sensitive to HOSCN inhibition than wild-type cells (12), a phenotype that is easiest to visualize at lower HOSCN concentrations than the 400 µM treatment used in **Figs 1**, **2**, and **4**. The concentration of HOSCN used did, however, impact the ability of NiSO_4_ to enhance HOSCN toxicity. When exposed to 200 µM HOSCN, wild-type *E. coli* were only modestly sensitized by addition of 300 µM NiSO_4_, unlike the Δ*rclA* mutant, which was sensitized quite strongly (**Fig 5**). This indicates that the mechanism by which Ni^2+^ enhances HOSCN toxicity cannot possibly depend on inhibition of RclA. The same was true of the Δ*rclABC* triple mutant (**Fig 5**), which additionally lacks the RclB and RclC proteins (49, 56), indicating that neither RclB nor RclC were the target of Ni^2+^ either. RclB and RclC protect *E. coli* against HOSCN (12) by currently unknown mechanisms, and while the Δ*rclABC* mutant was not more sensitive to inhibition by 200 µM HOSCN than the Δ*rclA* mutant under these growth conditions (**Fig 5**), it was completely inhibited by 300 µM HOSCN (**Supplemental Fig S5**), unlike the Δ*rclA* mutant, which was able to eventually recover growth even at 400 µM HOSCN (**Fig 4**).

**FIG 5.**
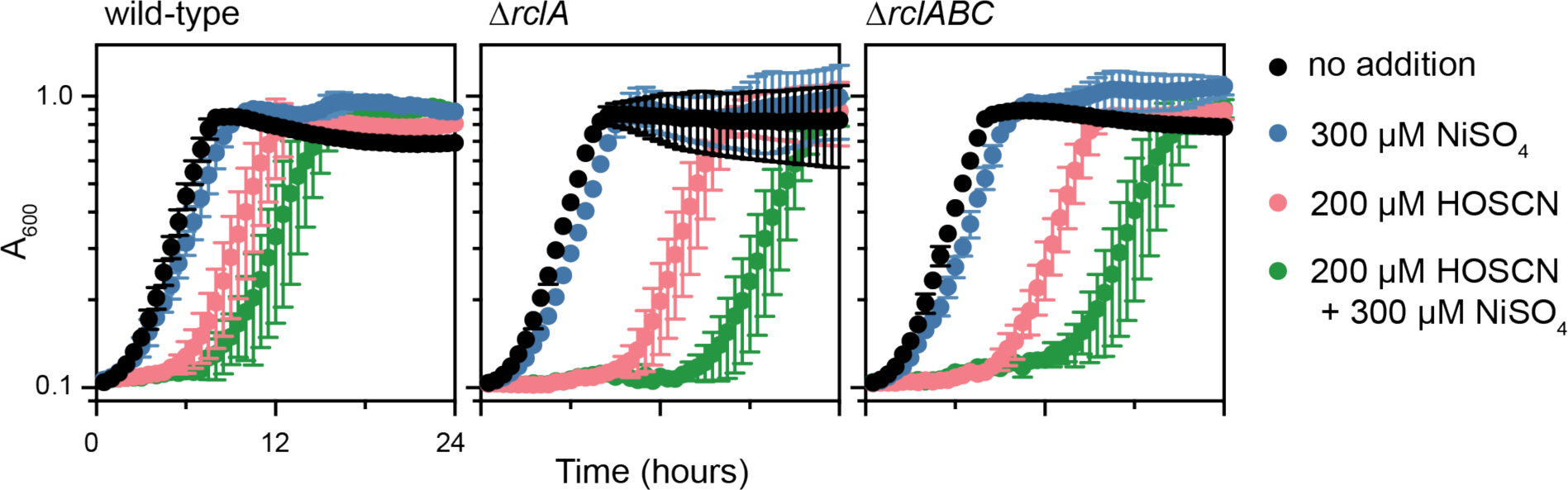
Nickel enhancement of HOSCN toxicity is independent of RclABC. *E. coli* strains MG1655 (wild-type), MJG1958 (MG1655 &*rclA747*), and MJG0901 (MG1655 &*rclABC1*) were grown in MOPS minimal medium without micronutrients containing 0.2% glucose, supplemented with 300 µM NiSO_4_ and/or 200 µM HOSCN, as indicated, incubating at 37°C with shaking and measuring A_600_ at 30-minute intervals for 24 hours (n=3 experimental replicates, with 3 technical replicates per experiment; error bars of 1 standard deviation).

### Glutathione is a major HOSCN defense in *E. coli*

Having eliminated the known HOSCN-specific defenses of *E. coli* (*i.e.* RclABC) as targets of Ni^2+^, I next moved on to testing whether other genes involved in related oxidative stress responses might be involved. Deletion of the *nemA*, *hslO*, and *mgsA* genes, encoding proteins involved in defense against HOCl in *E. coli* (41, 57, 58), had no impact on HOSCN resistance (**Fig 6A**). Similarly, while deletion of the genes encoding thioredoxin reductase (*trxB*)(20, 38) or either of the two thioredoxin homologs of *E. coli* (*trxA* and *trxC*)(20, 38, 59) modestly slowed the growth of *E. coli* under non-stress conditions, none of those mutants had substantial HOSCN sensitivity phenotypes (**Fig 6B**). The same was true of a Δ*grxA* mutant (**Fig 6C**) lacking glutaredoxin (21, 38) However, a Δ*gor* mutant lacking glutathione oxidoreductase (Gor)(38, 60) was extremely sensitive to HOSCN. The *E. coli* Δ*gor* strain was completely inhibited by 400 µM HOSCN (**Fig 6C**) and both Δ*gor* and Δ*gshA* mutants (which are unable to synthesize glutathione)(38, 61) were strongly inhibited by 100 µM HOSCN, a concentration which barely effected the growth of the wild-type (**Fig 7**). Glutathione is abundant in *E. coli* (62), is oxidized efficiently by HOSCN (63), and *S. pneumoniae gor* mutants are also very sensitive to HOSCN (19), so this phenotype was not an enormous surprise. The question, though, was whether Gor or glutathione are involved in the Ni^2+^-sensitization phenotype. Although the variability in the recovery time of HOSCN-stressed Δ*gor* and Δ*gshA* mutants was relatively high (note the large error bars in **Fig 7**), neither mutant appeared to be notably further sensitized to HOSCN by 300 µM NiSO_4_.

**FIG 6.**
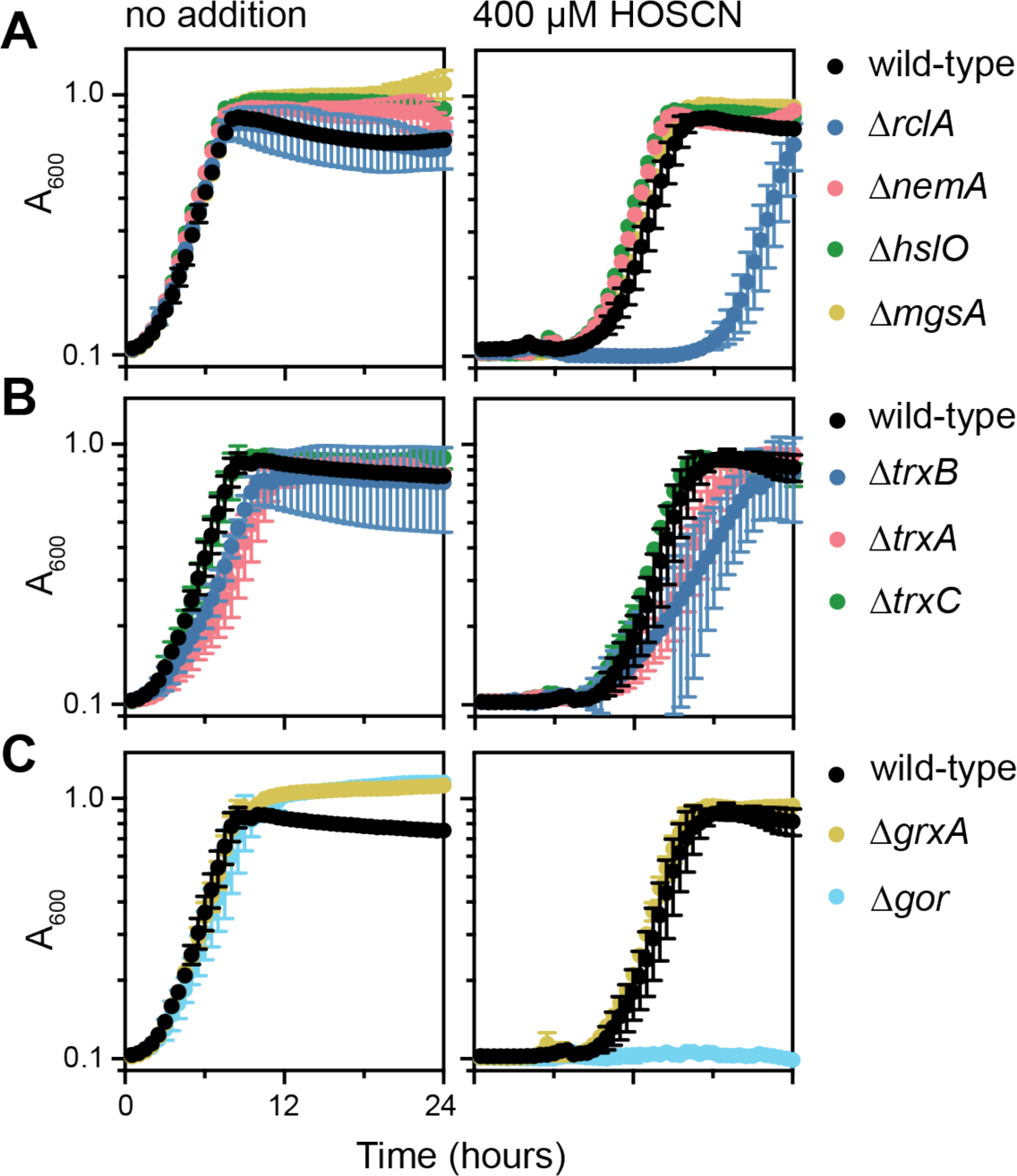
Glutathione reductase contributes to *E. coli* HOSCN resistance. *E. coli* strains MG1655 (wild-type), MJG0044 (MG1655 &*nemA728*), MJG0081 (MG1655 *hslO*::*kan*^+^), MJG0176 (MG1655 &*mgsA778*), MJG1958 (MG1655 &*rclA747*), MJG2371 (MG1655 &*trxB786*::*kan*^+^), MJG2385 (MG1655 &*trxA732*::*kan*^+^), MJG2387 (MG1655 &*trxC750*::*kan*^+^), MJG2388 (MG1655 &*grxA750*::*kan*^+^), MJG2398 (MG1655 &*gor-756*::*kan*^+^) were grown in MOPS minimal medium without micronutrients containing 0.2% glucose, with or without 400 µM HOSCN, as indicated, incubating at 37°C with shaking and measuring A_600_ at 30-minute intervals for 24 hours (n=3 experimental replicates, with 3-4 technical replicates per experiment; error bars of 1 standard deviation).

**FIG 7.**
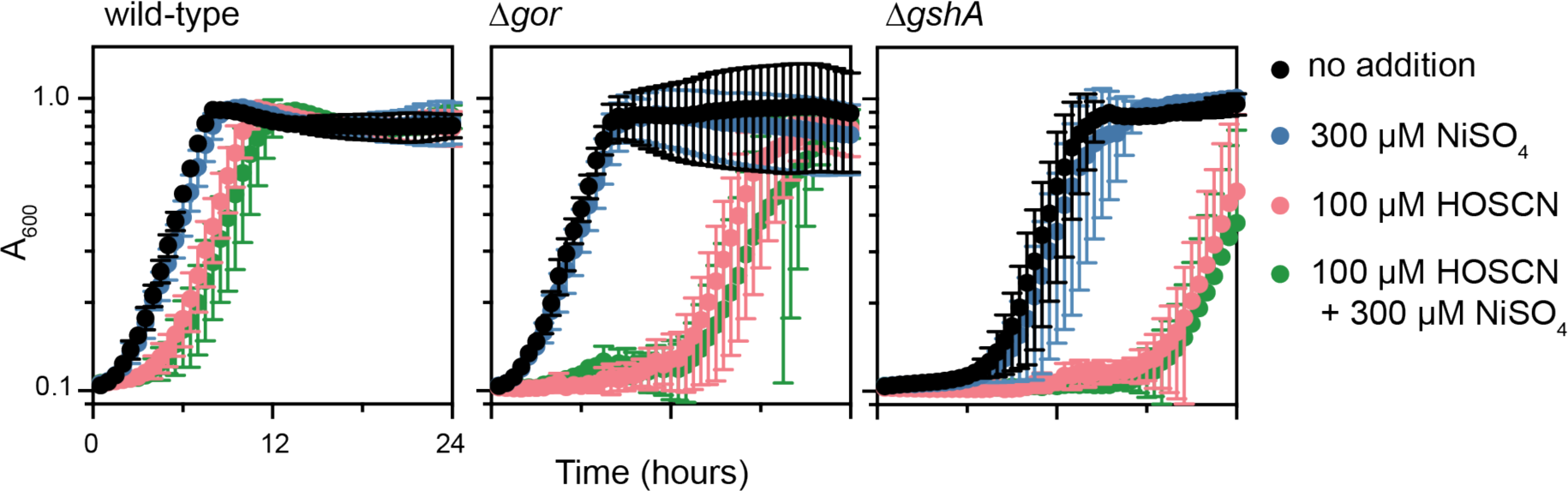
Glutathione mutants are extremely sensitive to HOSCN. *E. coli* strains MG1655 (wild-type), MJG2398 (MG1655 &*gor-756*::*kan*^+^), and MJG2485 (MG1655 &*gshA769*:*kan*^+^) were grown in MOPS minimal medium without micronutrients containing 0.2% glucose, with or without 100 µM HOSCN and/or 300 µM NiSO_4_, as indicated, incubating at 37°C with shaking and measuring A_600_ at 30-minute intervals for 24 hours (n=3 experimental replicates, with 4 technical replicates per experiment; error bars of 1 standard deviation).

### Glutathione reductase is inhibited by nickel, but not by HOSCN

The *in vivo* results in **Fig 7** suggested that Gor might be the target of Ni^2+^ in HOSCN-stressed cells but were not conclusive. I therefore used cell-free lysates of *E. coli* to directly test the impact of both NiSO_4_ and HOSCN on Gor activity *in vitro* (64, 65). NiSO_4_ inhibited Gor (**Fig 8A**), but HOSCN did not (**Fig 8B**), and HOSCN-oxidized lysates were as sensitive to inhibition by NiSO_4_ as lysates which had not been exposed to HOSCN (**Fig 8B**). Since polyP could counteract the HOSCN-sensitizing effect of NiSO_4_ *in vivo* (**Fig 2**), I added purified polyP to lysates to see if it could also suppress Gor inhibition *in vitro*. As shown in **Fig 8C**, 5 mM polyP (concentration expressed in terms of individual phosphate units due to the heterogenous length of commercially-available polyP) significantly reduced the ability of 1 mM NiSO_4_ to inhibit Gor activity, presumably due to its ability to chelate divalent metal cations (66–68). Addition of 5 mM of the chelator EDTA completely eliminated NiSO_4_ inhibition of Gor activity, and 5 mM polyP had no effect on Gor activity in the absence of NiSO_4_ (**Fig 8C**). These results support the model that Gor is the target that Ni^2+^ affects to sensitize *E. coli* to HOSCN. In contrast to their effects on Gor activity, 5 mM NiSO_4_ did not inhibit thioredoxin reductase (TrxB) activity in lysates, but HOSCN completely inactivated it (**Fig 8D**), consistent with previous reports that *E. coli* TrxB is HOSCN-sensitive (4).

**FIG 8.**
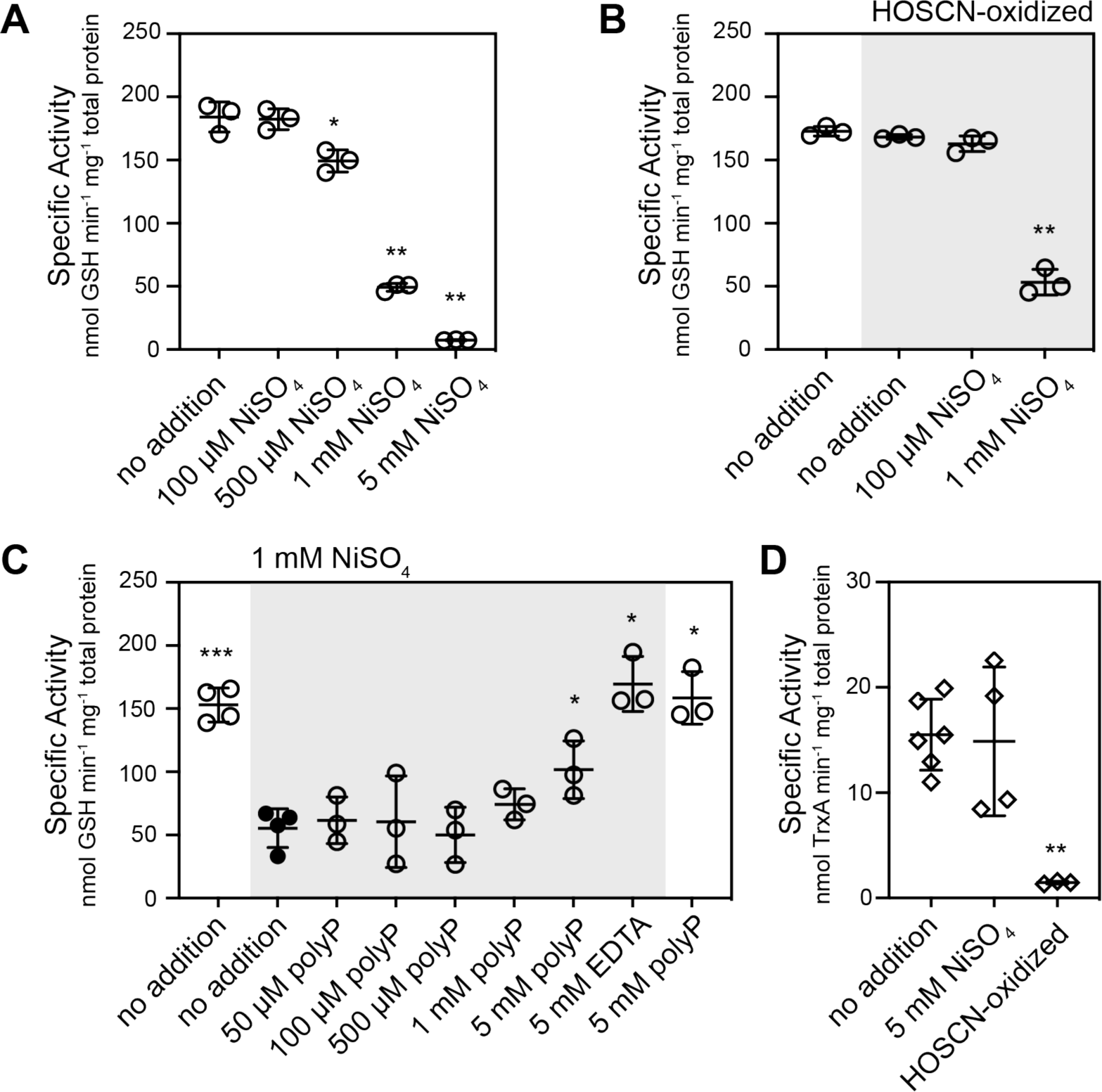
Glutathione reductase is inhibited by nickel but not by HOSCN. Glutathione and thioredoxin reductase activity was measured in cell-free lysates of *E. coli* MG1655. Glutathione reductase assays (**A**-**C**) contained 50 mM HEPES-KOH buffer (pH 8), 1.2 mM oxidized glutathione, 600 µM DTNB, 350 µM NADPH, and thioredoxin reductase assays (**D**) contained 50 mM HEPES-KOH buffer (pH 8), 500 nM *E. coli* thioredoxin 1 (TrxA), 0.1 mg ml^-1^ bovine serum albumin, 500 µM DTNB, and 240 µM NADPH. Reactions also contained the indicated concentrations of NiSO_4_, polyP, and/or EDTA, and were started by addition of cell lysate (1.63-16.3 µg total protein). DTNB oxidation to TNB was measured over time, using the TNB extinction coefficient at 412 nm of 14100 M^-1^ cm^-1^, then specific activities were calculated as nmol GSH or TrxA reduced min^-1^ mg^-1^ total protein (n=3-6 experimental replicates; error bars of 1 standard deviation). HOSCN-oxidized lysates (used as indicated in **B** and **D**) were prepared by incubating cell lysate (3.26 mg ml^-1^ total protein) for 5 min at room temperature with 3 mM HOSCN, then quenching unreacted HOSCN with an equal volume of 2 mM TNB. Asterisks indicate activities significantly different from that of (**A**, **B**, and **D**) lysate with no NiSO_4_ added or (**C**) lysate with 1 mM NiSO_4_ added (filled circles, one-way ANOVA with Holm-Sidak9s multiple comparisons test or mixed model with Dunnett9s multiple comparisons test)(* = P<0.05, ** = P<0.01, *** = P<0.001).

## DISCUSSION

The molecular details of the layered defenses that different bacteria employ to protect themselves against HOSCN are now beginning to be deciphered (11–19, 37), decades after HOSCN was first identified as an antimicrobial compound produced by the immune system (69). This paper now shows that glutathione is a key component in the *E. coli* HOSCN stress response, distinct from the protection provided by more HOSCN-specific defenses (*e.g.* RclABC)(12). HOSCN is a specific thiol oxidant (2) and glutathione is one of the most abundant metabolites in the *E. coli* cytoplasm (reaching a concentration of 17 mM in cells grown in minimal glucose medium)(62), so it is not surprising that HOSCN stress would result in glutathione oxidation, and indeed, low molecular weight thiols (*e.g.* glutathione and bacillithiol)(70, 71) appear to be important for defense against HOSCN in a variety of bacteria (19, 37). My results also explain a puzzling older result (35) by showing that Ni^2+^ inhibits *E. coli* Gor, thereby sensitizing *E. coli* to HOSCN. Ni^2+^ has also been reported to inhibit both yeast and bovine glutathione reductases (72, 73), so this may be a general property of these enzymes, and could potentially contribute to the oxidative stress observed when diverse organisms are exposed to toxic levels of nickel (74–76).

I was somewhat surprised to find that the thioredoxin system does not appear to play an important role in HOSCN response in *E. coli* (**Fig 6**), since *trxA* mutants of *S. pneumoniae* are sensitive to HOSCN (18) and TrxB’s sensitivity to HOSCN inactivation has been proposed to be partially responsible for the antimicrobial activity of HOSCN (4). Based on my results, this seems unlikely, at least for *E. coli*, although since *trxB gor* and *trxB gshA* double mutants are exceptionally sensitive to oxidative stress (36), concurrent inactivation of TrxB by HOSCN and Gor by Ni^2+^ or other metals (**Figs 3**, **8**) could amplify the individual impacts of these compounds and contribute to the Ni^2+^-sensitization phenotype (**Fig 1**).

One question that remains unanswered is whether the sensitivity of HOSCN defense enzymes to inhibition by metals (**Figs 3**, **8**) is relevant to the ability of bacteria to resist HOSCN produced by their mammalian hosts (11). Ni^2+^ is present in food and water at widely varying concentrations (76), and *E. coli* has genes encoding both Ni^2+^ import proteins (*e.g. nikABCDE*, *corA*)(38, 39, 77) and exporters (*e.g. rcnA*)(55), implying that it is exposed to varying Ni^2+^ concentrations in its natural environment in the large intestine, some of which can reach potentially toxic levels (74). Whether the 300-500 µM Ni^2+^ concentrations that sensitized *E. coli* to HOSCN in laboratory growth media occur under physiological conditions where *E. coli* is also exposed to HOSCN is difficult to determine. For one thing, to my knowledge no studies to date have attempted to quantify HOSCN in the large intestine, although the presence of dedicated HOSCN resistance genes (*e.g. rclABC*) in diverse intestinal bacteria suggests it is present in at least some situations (11, 49). Cu^2+^ and Zn^2+^ were potent inhibitors of RclA activity *in vitro* (**Fig 3**), and while solubility issues in the minimal media I used to grow *E. coli* prevented me from determining unambiguously whether either of these metals could inhibit RclA *in vivo*, that does not necessarily mean that it cannot happen in the host environment. Both Cu^2+^ and Zn^2+^ are concentrated in the phagosomes of immune cells as part of the antimicrobial arsenal of those cells (78) and can reach much higher concentrations than I could achieve here (*e.g.* > 100 µM Cu^2+^ in the phagosomes of macrophages that have phagocytosed *Mycobacterium* cells)(79). It is certainly possible that under those conditions, inhibition of RclA by Cu^2+^ and/or Zn^2+^ could have a physiologically relevant impact on the ability of bacteria to survive host-produced oxidative stress. The expression of the *rclABC* operon of *E. coli* is strongly induced upon phagocytosis by neutrophils (80) and *rclA* contributes to the ability of *Salmonella* to survive phagocytosis by macrophages (50), but more work will need to be done to determine whether and under what circumstances metals affect bacterial HOSCN defenses during interactions with the host immune system.

## MATERIALS AND METHODS

### Bacterial strains and growth conditions

All strains used in this study are listed in **Table 1**. I grew *E. coli* at 37°C in Lysogeny Broth (LB)(81) containing 5 g l^-1^ NaCl or in MOPS minimal medium (82) containing 2 g l^-1^ glucose, with kanamycin (25 or 50 µg ml^-1^) added to LB medium when appropriate. For *in vivo* experiments to assess the impact of metal ions on HOSCN resistance, I prepared MOPS medium without adding micronutrients (*i.e.* (NH_4_)_6_Mo_7_O_24_, H_3_BO_3_, CoCl_2_, CuSO_4_, MnCl_2_, ZnSO_4_, and CaCl_2_). Minimal medium was used for all *in* vivo tests of HOSCN sensitivity due to the known ability of the components of undefined rich media to react with antimicrobial oxidants, including HOSCN (83–85).

**TABLE 1.**
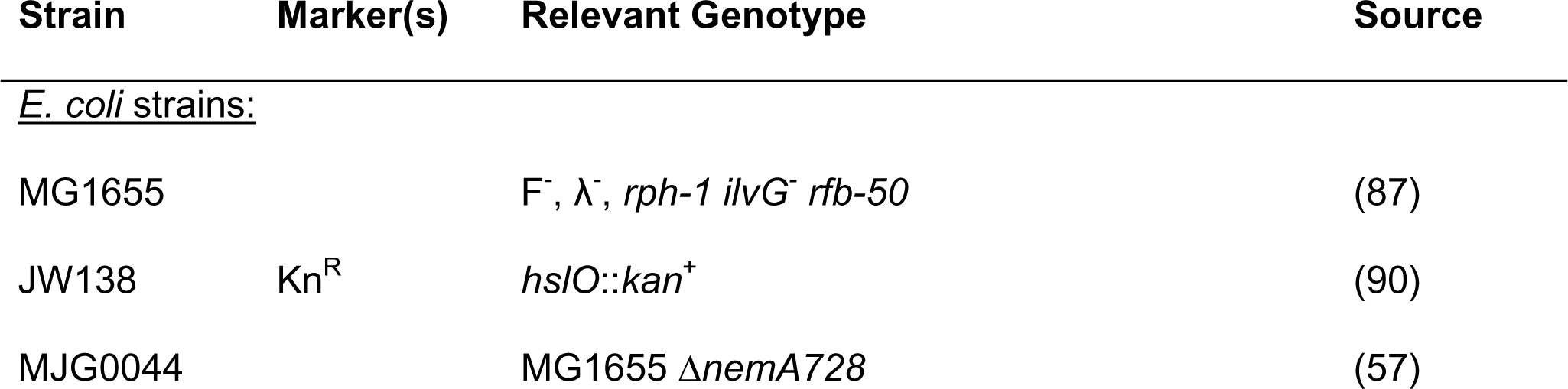
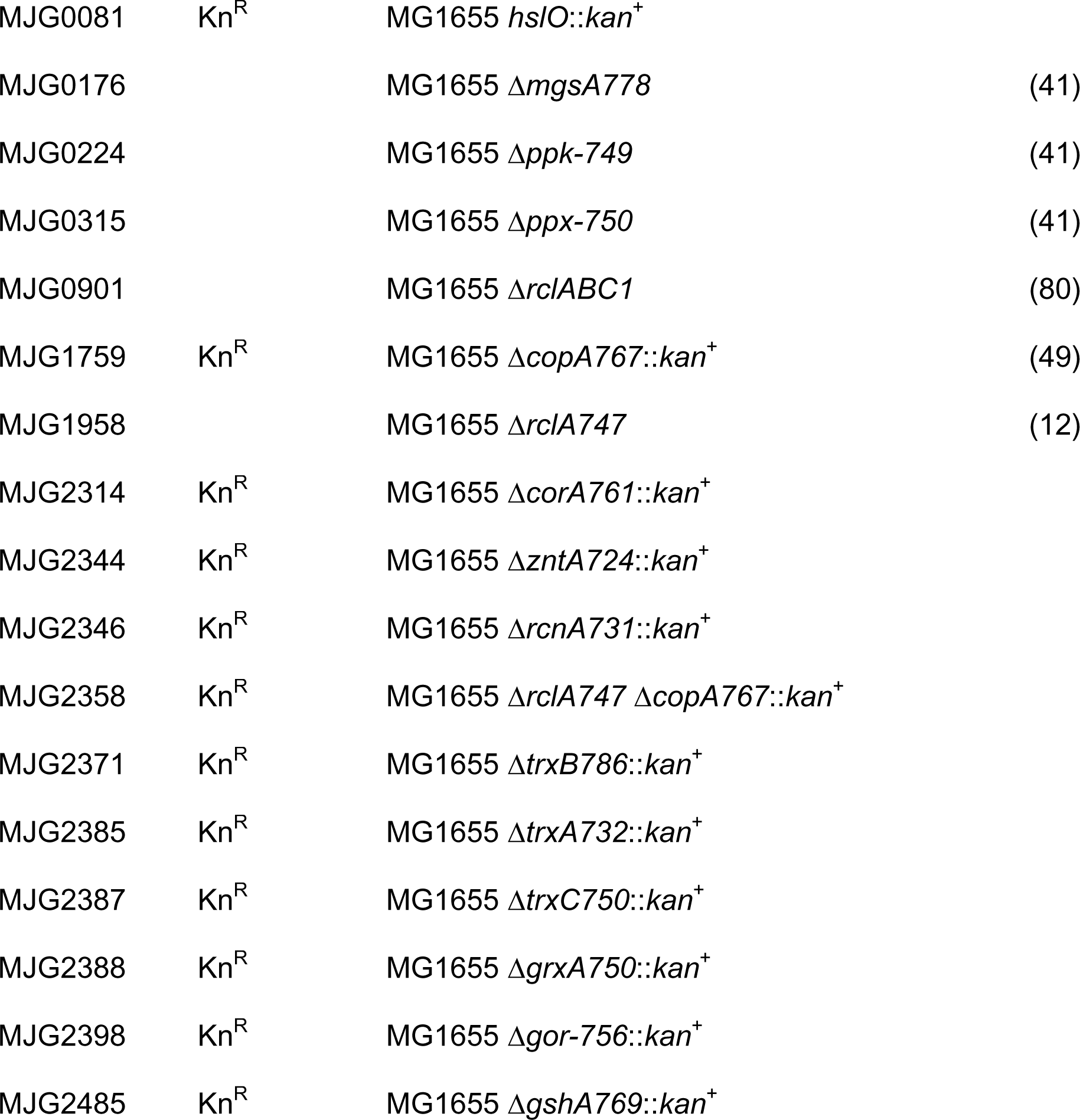
Strains used in this study. Unless otherwise indicated, strains were generated in the course of this work. Abbreviations: Kn^R^, kanamycin resistance.

| Strain | Marker(s) | Relevant Genotype | Source |
| --- | --- | --- | --- |
| <u>*E. coli* strains:</u> |  |  |  |
| MG1655 |  | F<sup>-</sup>, λ<sup>-</sup>, *rph-1 ilvG<sup>-</sup> rfb-50* | (87) |
| JW138 | Kn<sup>R</sup> | *hsIO::kan<sup>+</sup>* | (90) |
| MJG0044 |  | MG1655 Δ*nemA*728 | (57) |
| MJG0081 | Kn<sup>R</sup> | MG1655 *hslO::kan<sup>+</sup>* |  |
| MJG0176 | | MG1655 Δ*mgsA778* | (41) |
| MJG0224 | | MG1655 Δ*ppk-749* | (41) |
| MJG0315 | | MG1655 Δ*ppx-750* | (41) |
| MJG0901 | | MG1655 Δ*rclABC1* | (80) |
| MJG1759 | Kn<sup>R</sup> | MG1655 Δ*copA767::kan<sup>+</sup>* | (49) |
| MJG1958 | | MG1655 Δ*rclA747* | (12) |
| MJG2314 | Kn<sup>R</sup> | MG1655 Δ*corA761::kan<sup>+</sup>* | |
| MJG2344 | Kn<sup>R</sup> | MG1655 Δ*zntA724::kan<sup>+</sup>* | |
| MJG2346 | Kn<sup>R</sup> | MG1655 Δ*rcnA731::kan<sup>+</sup>* | |
| MJG2358 | Kn<sup>R</sup> | MG1655 Δ*rclA747* Δ*copA767::kan<sup>+</sup>* | |
| MJG2371 | Kn<sup>R</sup> | MG1655 Δ*trxB786::kan<sup>+</sup>* | |
| MJG2385 | Kn<sup>R</sup> | MG1655 Δ*trxA732::kan<sup>+</sup>* | |
| MJG2387 | Kn<sup>R</sup> | MG1655 Δ*trxC750::kan<sup>+</sup>* | |
| MJG2388 | Kn<sup>R</sup> | MG1655 Δ*grxA750::kan<sup>+</sup>* | |
| MJG2398 | Kn<sup>R</sup> | MG1655 Δ*gor-756::kan<sup>+</sup>* | |
| MJG2485 | Kn<sup>R</sup> | MG1655 Δ*gshA769::kan<sup>+</sup>* | |

### Databases and primer design

I obtained gene and protein sequences and other information from the Integrated Microbial Genomes database (86) and from EcoCyc (38). I designed PCR and sequencing primers with Web Primer (www.candidagenome.org/cgi-bin/compute/web-primer).

### Strain construction

All *E. coli* strains used in this study were derivatives of wild-type strain MG1655 (F^-^, λ^-^, *rph-1 ilvG^-^ rfb-50*) (87). I used P1*vir* phage transduction (88, 89) to move the *hslO*::*kan*^+^ allele from *E. coli* strain JW138 (90), which is a 737 bp internal deletion of *hslO* tagged with a kanamycin-resistance cassette (91), into MG1655, generating strain MJG0081 (MG1655 *hslO*::*kan*^+^). I used P1*vir* transduction to move the Δ*corA761*::*kan*^+^, Δ*copA767*::*kan*^+^, Δ*zntA724*::*kan*^+^, Δ*rcnA731*::*kan*^+^, Δ*trxB786*::*kan*^+^, Δ*trxA732*::*kan*^+^, Δ*trxC750*::*kan*^+^, Δ*grxA750*::*kan*^+^, Δ*gor-756*::*kan*^+^, and Δ*gshA769*::*kan*^+^ alleles from the Keio collection (92) into MG1655 or MJG1958 (MG1655 Δ*rclA747*)(12), generating strains MJG2314 (MG1655 Δ*corA761*::*kan*^+^), MJG2344 (MG1655 Δ*zntA724*::*kan*^+^), MJG2346 (MG1655 Δ*rcnA731*::*kan*^+^), MJG2358 (MG1655 Δ*rclA747* Δ*copA767*::*kan*^+^), MJG2371 (MG1655 Δ*trxB786*::*kan*^+^), MJG2385 (MG1655 Δ*trxA732*::*kan*^+^), MJG2387 (MG1655 Δ*trxC750*::*kan*^+^), MJG2388 (MG1655 Δ*grxA750*::*kan*^+^), MJG2398 (MG1655 Δ*gor-756*::*kan*^+^), and MJG2485 (MG1655 Δ*gshA769*::*kan*^+^). *E. coli corA* mutants are calcium-sensitive, so when phage were grown on *corA* strains, I used media containing only 5 mM CaCl_2_ and 15 mM MgSO_4_ (93). I confirmed chromosomal mutations by PCR and whole-genome sequencing (SeqCenter, Philadelphia, PA).

### Preparation of HOSCN

I prepared HOSCN with lactoperoxidase (LPO; Worthington Biochemical cat. # LS000151) as previously described (12). I combined 1.2 µM LPO and 7.5 mM NaSCN in 10 mM HEPES-KOH, pH 7.4 (for *in vitro* experiments) or MOPS minimal medium (82) containing 0.2% glucose but lacking micronutrients (for *in vivo* experiments), then added 0.8 mM H_2_O_2_ three times at one-minute intervals and incubated on ice for 10 minutes. I added 30 μg of catalase to degrade excess H_2_O_2_, then removed the proteins with a 10K MWCO protein concentrator (Millipore) and quantified the resulting HOSCN solution (typically 3.5 to 4 mM) with 5-thio-2-nitrobenzoic acid (TNB; extinction coefficient at 412 nm = 14100 M^-1^ cm^-1^) by measuring the loss of absorbance at 412 nm after adding HOSCN (63). Cu, but neither Zn nor Ni, was able to catalyze a small amount of TNB oxidation (**Fig S6**), which was taken into account when necessary.

### *In vivo* HOSCN stress treatment assays

I grew *E. coli* strains overnight at 37°C with shaking in MOPS minimal medium containing 0.2% glucose then normalized the cultures to A_600_ = 1 and rinsed twice with sterile MOPS minimal medium containing 0.2% glucose but lacking micronutrients. For experiments involving Δ*ppk* mutants (*i.e.* **Fig 2**), which grow very poorly in the absence of amino acids (47, 94), the resuspension solution also included 4% (w/v) casamino acids (Fisher Scientific cat. #BP1424). I diluted the resulting cell suspensions 1:40 into MOPS minimal medium without micronutrients containing 0.2% glucose, supplemented with HOSCN and/or metal salts, as indicated. I performed growth curves in 200-µl volumes in clear 96-well plates in a Tecan Spark plate reader, incubating at 37°C with shaking and measuring A_600_ at 30-minute intervals for 24 hours.

### *In vitro* HOSCN reductase activity assay

The purification of Strep-tagged RclA has been previously described (12, 49). I measured the HOSCN reductase activity of purified RclA aerobically in 10 mM HEPES-KOH buffer (pH 7.4) at 20°C. Each reaction (500 µl) contained 160 µM NADH and, where indicated, 100 µM HOSCN and the indicated concentrations of metal salts and was started by addition of 10 nM RclA. I quantified NADH consumption over time using a GENESYS 10S UV-Vis spectrophotometer (ThermoFisher), measuring A_340_ and using the NADH extinction coefficient at 340 nm of 6220 M^-1^ cm^-1^ to calculate NADH concentrations, then fit slopes to the resulting linear plots and calculated specific activities as µmol NADH consumed min^-1^ mg^-1^ RclA.

### Glutathione and thioredoxin reductase activity assays

I prepared cell-free lysates of *E. coli* MG1655 from 400-ml overnight cultures grown in LB as follows: I rinsed pelleted cells with 50 mM HEPES-KOH buffer (pH 8) and stored them at −80°C before thawing on ice and resuspending in 5 ml of the same buffer. I then lysed the cells by sonication on ice in a Fisher Model 120 Sonic Dismembrator (5 min @ 50% amplitude, 5 sec on, 5 sec off), removed debris by centrifugation (40 min @ 21,000 g @ 4°C), and dialyzed thoroughly into 50 mM HEPES-KOH buffer (pH 8) at 4°C using 12-14 kDa MWCO dialysis tubing before storing aliquots at −80°C. I measured the total protein content of the lysates using the Bradford assay (ThermoFisher). Where indicated, I oxidized lysates before use by diluting them 1:5 into buffer containing HOSCN (3 mM final concentration), incubating for 5 min at room temperature, then adding an equal volume of 2 mM TNB to quench unreacted HOSCN (4).

I measured glutathione reductase activity in *E. coli* lysates by a modification of a previously published method (64). Reactions (500 µl) contained 50 mM HEPES-KOH buffer (pH 8), 1.2 mM oxidized glutathione (GSSG; Fisher Scientific cat. # AAJ6371503), 600 µM 5,5’-dithiobis-(2-nitrobenzoic acid)(DTNB), 350 µM NADPH, and the indicated concentrations of NiSO_4_ and/or polyP (Acros Organics cat. # 390932500), and were started by addition of cell lysate (16.3 µg total protein). PolyP concentrations are expressed in terms of phosphate monomers, due to the heterogeneity in chain length of commercial polyP. I measured thioredoxin reductase activity in lysates by a modification of a similar previously published method (65). Those reactions (500 µl) contained 50 mM HEPES-KOH buffer (pH 8), 500 nM *E. coli* thioredoxin 1 (TrxA; Sigma-Aldrich cat. # T0910), 0.1 mg ml^-1^ bovine serum albumin, 500 µM DTNB, 240 µM NADPH, and the indicated concentrations of NiSO_4_ and were also started by addition of cell lysate (1.63-16.3 µg total protein). For both assays, I quantified DTNB oxidation to TNB over time using a GENESYS 10S UV-Vis spectrophotometer (ThermoFisher), measuring A_412_ and using the TNB extinction coefficient at 412 nm of 14100 M^-1^ cm^-1^ to calculate TNB concentrations, then fit slopes to the resulting linear plots and calculated specific activities as nmol GSSG or thioredoxin reduced min^-1^ mg^-1^ total protein.

### Statistical analyses

I used GraphPad Prism version 10.2.0 for Macintosh (GraphPad Software) to perform all statistical analyses, linear regressions, and graph generation. I based the graph color schemes on the color blindness-friendly “*Bright* qualitative color scheme” recommended of Dr. Paul Tol (95).

### Data availability

All strains generated in the course of this work are available from the author upon request. I deposited DNA sequencing data in the NIH Sequence Read Archive (accession # PRJNA943195), and all other raw data is available on FigShare (DOI: 10.6084/m9.figshare.c.7106581).

## ACKNOWLEDGEMENTS

This project was supported by University of Alabama at Birmingham Department of Microbiology development funds and NIH grant R35 GM124590. The author has no conflicts of interest to declare. Thanks to Dr. Ursula Jakob (U Michigan) for strain JW138, to Dr. Rhea Derke (UAB; currently at the Florida Department of Health – Duval County) for purified RclA, and to Julia Meredith (UAB) for helpful comments on the manuscript.

## REFERENCES

1. Arnhold J, Malle E. 2022. Halogenation Activity of Mammalian Heme Peroxidases. Antioxidants (Basel) 11.

2. Ashby MT. 2012. Chapter 8 - Hypothiocyanite, p 263-303. *In* Eldik Rv, Ivanović-Burmazović I (ed), Advances in Inorganic Chemistry: Inorganic/Bioinorganic Reaction Mechanisms, vol 64. Elsevier Inc.

3. Winterbourn CC, Kettle AJ, Hampton MB. 2016. Reactive Oxygen Species and Neutrophil Function. Annu Rev Biochem 85:765–92.

4. Chandler JD, Nichols DP, Nick JA, Hondal RJ, Day BJ. 2013. Selective metabolism of hypothiocyanous acid by mammalian thioredoxin reductase promotes lung innate immunity and antioxidant defense. J Biol Chem 288:18421–8.

5. Carlsson J, Iwami Y, Yamada T. 1983. Hydrogen peroxide excretion by oral streptococci and effect of lactoperoxidase-thiocyanate-hydrogen peroxide. Infect Immun 40:70–80.

6. Oram JD, Reiter B. 1966. The inhibition of streptococci by lactoperoxidase, thiocyanate and hydrogen peroxide. The effect of the inhibitory system on susceptible and resistant strains of group N streptococci. Biochem J 100:373–81.

7. Mickelson MN. 1966. Effect of lactoperoxidase and thiocyanate on the growth of Streptococcus pyogenes and Streptococcus agalactiae in a chemically defined culture medium. J Gen Microbiol 43:31–43.

8. Barrett TJ, Hawkins CL. 2012. Hypothiocyanous acid: benign or deadly? Chem Res Toxicol 25:263–73.

9. Chandler JD, Day BJ. 2015. Biochemical mechanisms and therapeutic potential of pseudohalide thiocyanate in human health. Free Radic Res 49:695–710.

10. San Gabriel PT, Liu Y, Schroder AL, Zoellner H, Chami B. 2020. The Role of Thiocyanate in Modulating Myeloperoxidase Activity during Disease. Int J Mol Sci 21.

11. Meredith JD, Gray MJ. 2023. Hypothiocyanite and host-microbe interactions. Mol Microbiol 119:302–311.

12. Meredith JD, Chapman I, Ulrich K, Sebastian C, Stull F, Gray MJ. 2022. Escherichia coli RclA is a highly active hypothiocyanite reductase. Proc Natl Acad Sci U S A 119:e2119368119.

13. Shearer HL, Loi VV, Weiland P, Bange G, Altegoer F, Hampton MB, Antelmann H, Dickerhof N. 2023. MerA functions as a hypothiocyanous acid reductase and defense mechanism in Staphylococcus aureus. Mol Microbiol 119:456–470.

14. Shearer HL, Pace PE, Paton JC, Hampton MB, Dickerhof N. 2022. A newly identified flavoprotein disulfide reductase Har protects Streptococcus pneumoniae against hypothiocyanous acid. J Biol Chem doi:10.1016/j.jbc.2022.102359:102359.

15. Farrant KV, Spiga L, Davies JC, Williams HD. 2020. Response of Pseudomonas aeruginosa to the Innate Immune System-Derived Oxidants Hypochlorous Acid and Hypothiocyanous Acid. J Bacteriol 203.

16. Groitl B, Dahl JU, Schroeder JW, Jakob U. 2017. Pseudomonas aeruginosa defense systems against microbicidal oxidants. Mol Microbiol 106:335–350.

17. Shearer HL, Kaldor CD, Hua H, Kettle AJ, Parker HA, Hampton MB. 2022. Resistance of Streptococcus pneumoniae to Hypothiocyanous Acid Generated by Host Peroxidases. Infect Immun 90:e0053021.

18. Shearer HL, Pace PE, Smith LM, Fineran PC, Matthews AJ, Camilli A, Dickerhof N, Hampton MB. 2023. Identification of Streptococcus pneumoniae genes associated with hypothiocyanous acid tolerance through genome-wide screening. J Bacteriol 205:e0020823.

19. Shearer HL, Paton JC, Hampton MB, Dickerhof N. 2022. Glutathione utilization protects Streptococcus pneumoniae against lactoperoxidase-derived hypothiocyanous acid. Free Radic Biol Med 179:24–33.

20. Lu J, Holmgren A. 2014. The thioredoxin antioxidant system. Free Radic Biol Med 66:75–87.

21. Meyer Y, Buchanan BB, Vignols F, Reichheld JP. 2009. Thioredoxins and glutaredoxins: unifying elements in redox biology. Annu Rev Genet 43:335–67.

22. Ashby MT, Kreth J, Soundarajan M, Sivuilu LS. 2009. Influence of a model human defensive peroxidase system on oral streptococcal antagonism. Microbiology (Reading) 155:3691–3700.

23. Lumikari M, Soukka T, Nurmio S, Tenovuo J. 1991. Inhibition of the growth of Streptococcus mutans, Streptococcus sobrinus and Lactobacillus casei by oral peroxidase systems in human saliva. Arch Oral Biol 36:155–60.

24. Thomas EL, Aune TM. 1978. Lactoperoxidase, peroxide, thiocyanate antimicrobial system: correlation of sulfhydryl oxidation with antimicrobial action. Infect Immun 20:456–63.

25. Thomas EL, Milligan TW, Joyner RE, Jefferson MM. 1994. Antibacterial activity of hydrogen peroxide and the lactoperoxidase-hydrogen peroxide-thiocyanate system against oral streptococci. Infect Immun 62:529–35.

26. Arnhold J. 2021. Heme Peroxidases at Unperturbed and Inflamed Mucous Surfaces. Antioxidants (Basel) 10.

27. Cupp-Sutton K, Ashby MT. 2021. Reverse Ordered Sequential Mechanism for Lactoperoxidase with Inhibition by Hydrogen Peroxide. Antioxidants (Basel) 10.

28. Seidel A, Parker H, Turner R, Dickerhof N, Khalilova IS, Wilbanks SM, Kettle AJ, Jameson GN. 2014. Uric acid and thiocyanate as competing substrates of lactoperoxidase. J Biol Chem 289:21937–49.

29. Sen A, Imlay JA. 2021. How Microbes Defend Themselves From Incoming Hydrogen Peroxide. Front Immunol 12:667343.

30. Garcia-Graells C, Van Opstal I, Vanmuysen SC, Michiels CW. 2003. The lactoperoxidase system increases efficacy of high-pressure inactivation of foodborne bacteria. Int J Food Microbiol 81:211–21.

31. Van Opstal I, Vanmuysen SC, Michiels CW. 2003. High sucrose concentration protects E. coli against high pressure inactivation but not against high pressure sensitization to the lactoperoxidase system. Int J Food Microbiol 88:1–9.

32. De Spiegeleer P, Sermon J, Vanoirbeek K, Aertsen A, Michiels CW. 2005. Role of porins in sensitivity of Escherichia coli to antibacterial activity of the lactoperoxidase enzyme system. Appl Environ Microbiol 71:3512–8.

33. Sermon J, Vanoirbeek K, De Spiegeleer P, Van Houdt R, Aertsen A, Michiels CW. 2005. Unique stress response to the lactoperoxidase-thiocyanate enzyme system in Escherichia coli. Res Microbiol 156:225–32.

34. De Spiegeleer P, Vanoirbeek K, Lietaert A, Sermon J, Aertsen A, Michiels CW. 2005. Investigation into the resistance of lactoperoxidase tolerant Escherichia coli mutants to different forms of oxidative stress. FEMS Microbiol Lett 252:315–9.

35. Sermon J, Wevers EM, Jansen L, De Spiegeleer P, Vanoirbeek K, Aertsen A, Michiels CW. 2005. CorA affects tolerance of Escherichia coli and Salmonella enterica serovar Typhimurium to the lactoperoxidase enzyme system but not to other forms of oxidative stress. Appl Environ Microbiol 71:6515–23.

36. Prinz WA, Aslund F, Holmgren A, Beckwith J. 1997. The role of the thioredoxin and glutaredoxin pathways in reducing protein disulfide bonds in the Escherichia coli cytoplasm. J Biol Chem 272:15661–7.

37. Loi VV, Busche T, Schnaufer F, Kalinowski J, Antelmann H. 2023. The neutrophil oxidant hypothiocyanous acid causes a thiol-specific stress response and an oxidative shift of the bacillithiol redox potential in Staphylococcus aureus. Microbiol Spectr 11:e0325223.

38. Karp PD, Paley S, Caspi R, Kothari A, Krummenacker M, Midford PE, Moore LR, Subhraveti P, Gama-Castro S, Tierrafria VH, Lara P, Muniz-Rascado L, Bonavides-Martinez C, Santos-Zavaleta A, Mackie A, Sun G, Ahn-Horst TA, Choi H, Covert MW, Collado-Vides J, Paulsen I. 2023. The EcoCyc Database (2023). EcoSal Plus 11:eesp00022023.

39. Snavely MD, Florer JB, Miller CG, Maguire ME. 1989. Magnesium transport in Salmonella typhimurium: 28Mg2+ transport by the CorA, MgtA, and MgtB systems. J Bacteriol 171:4761–6.

40. Seidel A. 2015. Kinetic Studies of Lactoperoxidase. Ph.D. University of Otago.

41. Gray MJ, Wholey WY, Wagner NO, Cremers CM, Mueller-Schickert A, Hock NT, Krieger AG, Smith EM, Bender RA, Bardwell JC, Jakob U. 2014. Polyphosphate is a primordial chaperone. Mol Cell 53:689–99.

42. Gray MJ, Jakob U. 2015. Oxidative stress protection by polyphosphate--new roles for an old player. Curr Opin Microbiol 24:1–6.

43. Bowlin MQ, Gray MJ. 2021. Inorganic polyphosphate in host and microbe biology. Trends Microbiol 29:1013–1023.

44. Fraley CD, Rashid MH, Lee SS, Gottschalk R, Harrison J, Wood PJ, Brown MR, Kornberg A. 2007. A polyphosphate kinase 1 (ppk1) mutant of Pseudomonas aeruginosa exhibits multiple ultrastructural and functional defects. Proc Natl Acad Sci U S A 104:3526–31.

45. Crooke E, Akiyama M, Rao NN, Kornberg A. 1994. Genetically altered levels of inorganic polyphosphate in Escherichia coli. J Biol Chem 269:6290–5.

46. Akiyama M, Crooke E, Kornberg A. 1993. An exopolyphosphatase of Escherichia coli. The enzyme and its ppx gene in a polyphosphate operon. J Biol Chem 268:633–9.

47. Kuroda A, Tanaka S, Ikeda T, Kato J, Takiguchi N, Ohtake H. 1999. Inorganic polyphosphate kinase is required to stimulate protein degradation and for adaptation to amino acid starvation in Escherichia coli. Proc Natl Acad Sci U S A 96:14264–9.

48. Gray MJ. 2020. Interactions between DksA and Stress-Responsive Alternative Sigma Factors Control Inorganic Polyphosphate Accumulation in Escherichia coli. J Bacteriol 202.

49. Derke RM, Barron AJ, Billiot CE, Chaple IF, Lapi SE, Broderick NA, Gray MJ. 2020. The Cu(II) Reductase RclA Protects Escherichia coli against the Combination of Hypochlorous Acid and Intracellular Copper. mBio 11.

50. Baek Y, Kim J, Ahn J, Jo I, Hong S, Ryu S, Ha NC. 2020. Structure and function of the hypochlorous acid-induced flavoprotein RclA from Escherichia coli. J Biol Chem 295:3202–3212.

51. Shin H, Baek Y, Lee D, Xu Y, Kwon Y, Jo I, Ha NC. 2023. Structural and Functional Analyses of the Flavoprotein Disulfide Reductase FN0820 of Fusobacterium nucleatum. J Microbiol 61:1033–1041.

52. Meydan S, Klepacki D, Karthikeyan S, Margus T, Thomas P, Jones JE, Khan Y, Briggs J, Dinman JD, Vazquez-Laslop N, Mankin AS. 2017. Programmed Ribosomal Frameshifting Generates a Copper Transporter and a Copper Chaperone from the Same Gene. Mol Cell 65:207–219.

53. Rensing C, Fan B, Sharma R, Mitra B, Rosen BP. 2000. CopA: An Escherichia coli Cu(I)-translocating P-type ATPase. Proc Natl Acad Sci U S A 97:652–6.

54. Rensing C, Mitra B, Rosen BP. 1997. The zntA gene of Escherichia coli encodes a Zn(II)-translocating P-type ATPase. Proc Natl Acad Sci U S A 94:14326–31.

55. Rodrigue A, Effantin G, Mandrand-Berthelot MA. 2005. Identification of rcnA (yohM), a nickel and cobalt resistance gene in Escherichia coli. J Bacteriol 187:2912–6.

56. Parker BW, Schwessinger EA, Jakob U, Gray MJ. 2013. The RclR protein is a reactive chlorine-specific transcription factor in Escherichia coli. J Biol Chem 288:32574–32584.

57. Gray MJ, Wholey WY, Parker BW, Kim M, Jakob U. 2013. NemR is a bleach-sensing transcription factor. J Biol Chem 288:13789–98.

58. Winter J, Ilbert M, Graf PC, Ozcelik D, Jakob U. 2008. Bleach activates a redox-regulated chaperone by oxidative protein unfolding. Cell 135:691–701.

59. Ritz D, Patel H, Doan B, Zheng M, Aslund F, Storz G, Beckwith J. 2000. Thioredoxin 2 is involved in the oxidative stress response in Escherichia coli. J Biol Chem 275:2505–12.

60. Perham RN. 1987. Glutathione reductase from Escherichia coli: mutation, cloning and sequence analysis of the gene. Biochem Soc Trans 15:730–3.

61. Greenberg JT, Demple B. 1986. Glutathione in Escherichia coli is dispensable for resistance to H2O2 and gamma radiation. J Bacteriol 168:1026–9.

62. Bennett BD, Kimball EH, Gao M, Osterhout R, Van Dien SJ, Rabinowitz JD. 2009. Absolute metabolite concentrations and implied enzyme active site occupancy in Escherichia coli. Nat Chem Biol 5:593–9.

63. Nagy P, Jameson GN, Winterbourn CC. 2009. Kinetics and mechanisms of the reaction of hypothiocyanous acid with 5-thio-2-nitrobenzoic acid and reduced glutathione. Chem Res Toxicol 22:1833–40.

64. Davis NK, Greer S, Jones-Mortimer MC, Perham RN. 1982. Isolation and mapping of glutathione reductase-negative mutants of Escherichia coli K12. J Gen Microbiol 128:1631–4.

65. Krause G, Holmgren A. 1991. Substitution of the conserved tryptophan 31 in Escherichia coli thioredoxin by site-directed mutagenesis and structure-function analysis. J Biol Chem 266:4056–66.

66. Beaufay F, Quarles E, Franz A, Katamanin O, Wholey WY, Jakob U. 2020. Polyphosphate Functions In Vivo as an Iron Chelator and Fenton Reaction Inhibitor. mBio 11.

67. Keasling JD. 1997. Regulation of intracellular toxic metals and other cations by hydrolysis of polyphosphate. Ann N Y Acad Sci 829:242–9.

68. Rudat AK, Pokhrel A, Green TJ, Gray MJ. 2018. Mutations in Escherichia coli Polyphosphate Kinase That Lead to Dramatically Increased In Vivo Polyphosphate Levels. J Bacteriol 200.

69. Hoogendoorn H, Piessens JP, Scholtes W, Stoddard LA. 1977. Hypothiocyanite ion; the inhibitor formed by the system lactoperoxidase-thiocyanate-hydrogen peroxide. I. Identification of the inhibiting compound. Caries Res 11:77–84.

70. Dumitrescu DG, Hatzios SK. 2023. Emerging roles of low-molecular-weight thiols at the host-microbe interface. Curr Opin Chem Biol 75:102322.

71. Van Laer K, Hamilton CJ, Messens J. 2013. Low-molecular-weight thiols in thiol-disulfide exchange. Antioxid Redox Signal 18:1642–53.

72. Tandogan B, Ulusu NN. 2007. The inhibition kinetics of yeast glutathione reductase by some metal ions. J Enzyme Inhib Med Chem 22:489–95.

73. Tandogan B, Ulusu NN. 2010. Inhibition of purified bovine liver glutathione reductase with some metal ions. J Enzyme Inhib Med Chem 25:68–73.

74. Macomber L, Hausinger RP. 2011. Mechanisms of nickel toxicity in microorganisms. Metallomics 3:1153–62.

75. Valko M, Morris H, Cronin MT. 2005. Metals, toxicity and oxidative stress. Curr Med Chem 12:1161–208.

76. Chain EPoCitF, Schrenk D, Bignami M, Bodin L, Chipman JK, Del Mazo J, Grasl-Kraupp B, Hogstrand C, Hoogenboom LR, Leblanc JC, Nebbia CS, Ntzani E, Petersen A, Sand S, Schwerdtle T, Vleminckx C, Wallace H, Guerin T, Massanyi P, Van Loveren H, Baert K, Gergelova P, Nielsen E. 2020. Update of the risk assessment of nickel in food and drinking water. EFSA J 18:e06268.

77. Eitinger T, Mandrand-Berthelot MA. 2000. Nickel transport systems in microorganisms. Arch Microbiol 173:1–9.

78. Sheldon JR, Skaar EP. 2019. Metals as phagocyte antimicrobial effectors. Curr Opin Immunol 60:1–9.

79. Wagner D, Maser J, Lai B, Cai Z, Barry CE, 3rd, Honer Zu Bentrup K, Russell DG, Bermudez LE. 2005. Elemental analysis of Mycobacterium avium-, Mycobacterium tuberculosis-, and Mycobacterium smegmatis-containing phagosomes indicates pathogen-induced microenvironments within the host cell’s endosomal system. J Immunol 174:1491–500.

80. Konigstorfer A, Ashby LV, Bollar GE, Billiot CE, Gray MJ, Jakob U, Hampton MB, Winterbourn CC. 2021. Induction of the reactive chlorine-responsive transcription factor RclR in Escherichia coli following ingestion by neutrophils. Pathog Dis 79.

81. Bertani G. 1951. Studies on lysogenesis. I. The mode of phage liberation by lysogenic Escherichia coli. J Bacteriol 62:293–300.

82. Neidhardt FC, Bloch PL, Smith DF. 1974. Culture medium for enterobacteria. J Bacteriol 119:736–47.

83. De Spiegeleer P, Sermon J, Lietaert A, Aertsen A, Michiels CW. 2004. Source of tryptone in growth medium affects oxidative stress resistance in Escherichia coli. J Appl Microbiol 97:124–33.

84. Ashby LV, Springer R, Hampton MB, Kettle AJ, Winterbourn CC. 2020. Evaluating the bactericidal action of hypochlorous acid in culture media. Free Radic Biol Med 159:119–124.

85. Verspecht T, Ghesquiere J, Bernaerts K, Boon N, Teughels W. 2021. Evaluating the intrinsic capacity of oral bacteria to produce hydrogen peroxide (H2O2) in liquid cultures: Interference by bacterial growth media. J Microbiol Methods 182:106170.

86. Markowitz VM, Chen IM, Palaniappan K, Chu K, Szeto E, Grechkin Y, Ratner A, Jacob B, Huang J, Williams P, Huntemann M, Anderson I, Mavromatis K, Ivanova NN, Kyrpides NC. 2012. IMG: the Integrated Microbial Genomes database and comparative analysis system. Nucleic Acids Res 40:D115–22.

87. Blattner FR, Plunkett G, 3rd, Bloch CA, Perna NT, Burland V, Riley M, Collado-Vides J, Glasner JD, Rode CK, Mayhew GF, Gregor J, Davis NW, Kirkpatrick HA, Goeden MA, Rose DJ, Mau B, Shao Y. 1997. The complete genome sequence of Escherichia coli K-12. Science 277:1453–62.

88. Silhavy TJ, Berman ML, Enquist LW (ed). 1984. Experiments with gene fusions. Cold Spring Harbor Laboratory, Cold Spring Harbor, NY.

89. Gray MJ. 2019. Inorganic Polyphosphate Accumulation in Escherichia coli Is Regulated by DksA but Not by (p)ppGpp. J Bacteriol 201:e00664–18.

90. Barth E, Gora KV, Gebendorfer KM, Settele F, Jakob U, Winter J. 2009. Interplay of cellular cAMP levels, sigmaS activity and oxidative stress resistance in Escherichia coli. Microbiology (Reading) 155:1680–1689.

91. Jakob U, Muse W, Eser M, Bardwell JC. 1999. Chaperone activity with a redox switch. Cell 96:341–52.

92. Baba T, Ara T, Hasegawa M, Takai Y, Okumura Y, Baba M, Datsenko KA, Tomita M, Wanner BL, Mori H. 2006. Construction of Escherichia coli K-12 in-frame, single-gene knockout mutants: the Keio collection. Mol Syst Biol 2:2006 0008.

93. Park MH, Wong BB, Lusk JE. 1976. Mutants in three genes affecting transport of magnesium in Escherichia coli: genetics and physiology. J Bacteriol 126:1096–103.

94. Gray MJ. 2019. Inorganic Polyphosphate Accumulation in Escherichia coli Is Regulated by DksA but Not by (p)ppGpp. J Bacteriol 201.

95. Tol P. 2021. “Color Schemes”; Technical note SRON/EPS/TN/09-002 3.2. SRON. https://personal.sron.nl/~pault/data/colourschemes.pdf. Accessed July 12, 2023.

